# Functional differences in electrolyte transport between the mouse proximal and distal trachea

**DOI:** 10.64898/2026.02.26.708204

**Authors:** Tábata Apablaza, Sandra Villanueva, Araceli Olave-Ruiz, Anita Guequen, Marcelo A. Catalán, Carlos A Flores

## Abstract

**Aim:** The mammalian tracheal epithelium is composed by different cell types unevenly distributed along the proximal-distal axis. Nevertheless, variations in expression and function of ion channels and transporters participating in fluid absorption and secretion had never been studied separately in proximal and distal sections of the mouse trachea. In this work, we aim to characterize basal and stimulated absorption and secretion of fluid obtained from proximal and distal trachea from the same animal.

**Methods:** Ussing chamber experiments were performed using a custom-made tissue slider that allowed the mounting small tracheal sections, where response to agonists and blockers was recorded. The role of the NKCC1 co-transporter was studied using the *Slc12a2*^-/-^ mouse. A genetically tomato-induced mouse model was used to assess co-expression of NKCC1 and ASCL3 by immunofluorescence. Animals were instilled with different interleukins (ILs) to determine changes in absorption, secretion and mucus properties.

**Results:** Proximal trachea didn’t participate in sodium absorption but exhibited higher cAMP- and succinate-induced anion secretion than the distal section. NBCe1-dependent bicarbonate and TMEM16A-driven chloride secretion was significantly higher in the distal section. NKCC1+ cells were found in the submucosal glands (SMGs) and abundant patches of NKCC1+ cells in the distal region. Isolated NKCC1+ cells co-expressing ASCL3 were also detected. ILs treatment changed the electrophysiological properties of the distal but not the proximal trachea.

**Conclusions:** Our experiments determined that the mouse trachea organizes its functions differentially in the proximal and distal sections, based in the functional distribution of channels, transporters and receptors. While the distal trachea drastically changed its responses to agonists inducing anion secretion the proximal trachea was unperturbed by the action of ILs.

## INTRODUCTION

The mammalian trachea is a cylindrical tube extending from the larynx downward its bifurcation into the left and right main bronchi (Proctor, 1977). It is covered by a layer of pseudostratified epithelium composed by different cell types that through the coordination of diverse functions including fluid absorption and secretion, mucus secretion and cilia beating, orchestrate the mucociliary clearance (MCC), a fundamental mechanism in airway and lung defense (Bustamante-Marin & Ostrowski, 2017; Davis & Wypych, 2021).

The impairment of ion transport in the airways results in MCC dysfunction and leads to severe lung function decay, as notably evident in cystic fibrosis (CF) and chronic obstructive pulmonary disease (COPD), where the reduction of CFTR-driven anion secretion lead to increased mucus viscosity, airway plugging and airflow obstructions (Dransfield *et al*., 2013; Saint-Criq & Gray, 2017). Similarly, mouse models with increased absorption of sodium or reduced secretion of bicarbonate exhibit a similar phenotype characterized by increased mucus viscosity, decreased MCC, mucus plugging, epithelial damage and inflammatory infiltration (Mall *et al*., 2004; Saint-Criq *et al*., 2022). Even though, the murine trachea has different features compared to the human, the mouse offers the possibility to use freshly excised tissue for electrophysiological recordings, allowing the study of the epithelium in their native environment and in connection with other cell types, like immune, neurons and smooth muscle, whose function can modulate or are modulated by the epithelium (Yang *et al*., 2017; O’Leary *et al*., 2019; Hewitt & Lloyd, 2021; Hollenhorst *et al*., 2022).

Another advantage of fresh tissue is that maintains the native distribution of different cell types and the anatomical differences along the proximal-distal axis of the mouse trachea. For example, submucosal glands (SMGs) are exclusively found in the most proximal region. Furthermore basal, serous and goblet cells distribute differentially along this axis. These regional variations suggest that epithelial functions, including ion transport, may also differ along the length of the trachea (Pack *et al*., 1980; Rock *et al*., 2010; Mahmoud *et al*., 2022; Peter Di & Mou, 2024). Nevertheless, the electrophysiological features of proximal and distal mouse (and other species) tracheal epithelium have never been studied individually.

Previously we observed that in some Ussing chamber experiments there was no response to the epithelial Na^+^-channel (ENaC) blocker amiloride, thus several experiments were not considered in the analysis or were discarded due to supposed damage during the dissection, even though, the observed transepithelial resistance (R_TE_) indicated an intact epithelial layer (Gianotti *et al*., 2016; Vega *et al*., 2020). This variability on amiloride responses is also addressed by other group (Grubb *et al*., 2001), thus we wondered if the difficulty to determine amiloride-sensitive sodium absorption was due to differences in the mounting of the tissue in the tissue sliders of the Ussing chambers, more specifically whether the proximal or the distal part of the trachea was left exposed during the experiment. To address this issue and to study regional differences we used specially designed tissue sliders to perform electrophysiological experiments on the proximal and distal parts of the same mouse trachea. Our results demonstrate that proximal and distal regions of the trachea bear different sodium absorption capacity, different responses to agonists for anion secretion, and a differential contribution of TMEM16A and NKCC1 driven chloride secretion and NBCe1-depdendent bicarbonate secretion. We also observed that the distal trachea can differentially modify ion absorption and secretion in response to different interleukins (ILs), while the proximal trachea remained unchanged.

## METHODS

All the material submitted is conform with good publishing practice in physiology (Jensen & Persson, 2022).

### Animals

Mice were housed at the Centro de Estudios Cientificos (CECs) animal facility. Wild type, *R26^tdTOMATO^* (The Jackson Laboratories, US) and *Ascl3*^P2A-GCE^ strain (Uchida *et al*., 2025)(kindly donated by Dr Catherine Ovitt, University of Rochester) were bred on the C57BL/6J background, but *Slc12a2* knockout mice which were on a Black Swiss/129 SvJ hybrid background (Flagella *et al*., 1999). Mice aged 2–5 months old were used.

### Ethics statement

All experimental procedures were approved by CECs Institutional Animal Care and Use Committee (#2022–04). The study was carried out in compliance with the ARRIVE guidelines.

### Reagents

Amiloride hydrochloride hydrate (A7410), uridine 5′-triphosphate trisodium salt hydrate (UTP) (U6750), Forskolin (FSK) (F6886), ANI9 (SML1813), S0859 (SML0638), Succinate acid disodium salt (224731) and Carbachol (C4382) were obtained from Sigma-Aldrich. Recombinant mouse interleukins were obtained from PeproTech® (Thermo Fisher Scientific), including IL-4 (Cat. No. 214-14), IL-1β (Cat. No. 211-11B), IL-17A (Cat. No. 210-17), and IL-13 (Cat. No. 210-13).

### Ussing chamber measurements

Mouse tracheae were divided into two sections: the proximal portion, from the cricoid cartilage to cartilage number 7, and the distal portion, dissected from cartilage number 7 to the main bronchi. Both sections were mounted in P2306CF tissue holders with an oval area of 0.02 cm² and placed in Ussing chambers (Physiologic Instruments Inc., San Diego, CA, USA). Tissues were bathed in a bicarbonate-buffered solution (adjusted to pH 7.4 with HCl) composed of (in mM): 120 NaCl, 25 NaHCO_₃_, 3.3 KH_₂_PO_₄_, 0.8 K_₂_HPO_₄_, 1.2 MgCl_₂_, and 1.2 CaCl_₂_. Additionally, 10 mM D-glucose was added to the basolateral side. Both solutions were continuously gassed with a mixture of 5% CO_₂_ and 95% O_₂_ and maintained at 37 °C. The transepithelial potential difference, referenced to the serosal side, was measured using a VCC MC2 amplifier (Physiologic Instruments Inc.). Equivalent short-circuit currents (Ieq) were calculated according to Ohm’s law. Sodium absorption via the epithelial sodium channel (ENaC) was inhibited with 10 µM amiloride. cAMP-dependent anion secretion was stimulated with 1 µM forskolin (FSK), while Ca²_⁺_-dependent anion secretion was induced with 300 µM succinate (Succ), 25 µM carbachol (CCh), and 10 µM UTP. ΔI_eq_ values were calculated as the amiloride-sensitive current (ΔAmiloride) and as the area under the curve (A.U.C.) of the first 2 minutes following the addition of FSK, Succ, CCh or UTP using the Acquire & Analyze software version 2.3. The apical chamber was washed 2 times with fresh buffer before the addition of CCh and UTP, and S0859 and ANI9 were also added before the agonists. Tissues with R_TE_ values below 50 Ω·cm² were excluded, as they were considered unsuitable for reliable electrophysiological measurements.

### Tissue procession and immunostaining

Wild-type (WT) and genetically modified mice were deeply anesthetized with isoflurane (Baxter), and euthanized by exsanguination via severing of the inferior vena cava. Animals were perfused with 10%/PBS, and tracheae and lungs were dissected and post-fixed overnight in 10% formalin/PBS at 4 °C. Tissues were paraffin-embedded and sectioned at 5 µm.

Sections were subjected to antigen retrieval in EDTA buffer (pH 8.0), blocked with 2.5% normal goat serum (Vector Laboratories, cat# S-1012), and incubated overnight at 4 °C with primary antibodies. After washing, sections were incubated for 2 h at room temperature with the appropriate Alexa Fluor–conjugated secondary antibodies (Invitrogen). Nuclear counterstaining was performed using either propidium iodide or DAPI, as indicated below.

### NKCC1 immunostaining in WT and *Nkcc1*-null mice

8-week-old WT mice were incubated with anti-NKCC1 antibody (D2O8R XP; Cell Signaling, cat# 85403; 1:100), followed by Alexa Fluor 488 goat anti-rabbit IgG (1:2000; Invitrogen, cat# A-11008). Nuclei were counterstained with propidium iodide (1:2000; Invitrogen, cat# P21493). To validate antibody specificity, tracheal tissue from *Slc12a2⁻ /⁻* (Nkcc1-null) mice was processed in parallel and used as a negative control.

### NKCC1 immunostaining in Ascl3 lineage-traced mice

5-month-old *Ascl3*^P2A-GCE^; *R26*^tdTomato^ mice received a single intraperitoneal injection of tamoxifen (Sigma-Aldrich, T5648; 0.2 mg/g body weight). Two weeks later, animals were perfused and processed as described above. Sections were incubated with anti-NKCC1 antibody (1:200) together with Living Colors® DsRed monoclonal antibody (Takara, cat# 632392), followed by Alexa Fluor 488 goat anti-rabbit IgG and Alexa Fluor 555 goat anti-mouse IgG (both 1:2000). Nuclei were counterstained with DAPI (1:2000; Invitrogen, cat# D3571).

### Image acquisition

NKCC1 immunostaining in WT and *Nkcc1*-null mice was imaged using a confocal microscope (Olympus FV1000). Lineage-tracing experiments were imaged using a Zeiss LSM 990 confocal microscope (Carl Zeiss Microscopy).

For LSM 990 acquisitions, three fluorescence channels were sequentially acquired using laser lines at 405 nm (0.9%), 488 nm (0.2%), and 543 nm (0.3%), with the pinhole set to 1.0 Airy unit for all channels. Large-area tracheal images were acquired using a tile scan consisting of 112 tiles with a Plan-Apochromat 20×/0.8 objective. Colocalization analysis was performed using Manders’ overlap coefficients with intensity-weighted correlation in ZEN 3.12 software (Zeiss). Manual thresholding was applied, and region-of-interest (ROI)-based analysis was conducted at the single-cell level.

### Interleukin instillation

Interleukins were prepared at a final concentration of 1 µg/mL in BSA 0.1% (vehicle). Each animal received a total dose of 30 ng of interleukin, administered in a final volume of 30 µL. Prior to instillation, mice were anesthetized with isoflurane (Baxter) using a laboratory animal induction chamber. Adequate depth of anesthesia was confirmed by the reduction of voluntary body movement, decreased whisker movement, and the presence of a regular breathing pattern. The animal was subsequently positioned at an approximately 70° vertical angle to optimize intranasal administration. 15 µL of the interleukin solution was delivered into each nostril using a micropipette. Following instillation, mice were placed on a heating pad until full recovery from anesthesia and returned to their cages. The experiments were performed 48h after instillation.

### Mucus transport and length measurements

For mucus transport we used the technique described previously (Apablaza *et al*., 2025). Briefly, dissected tissues were maintained in cold bicarbonate-buffered solution gassed with 5% CO_2_ and 95% O_2_. Tracheas was meticulously cleaned and mounted on a 35mm diameter plate coated with Sylgard 184 (The Dow Chemical Company). Lungs were pinned, and small incisions were made to release air, preventing the formation of bubbles that could interfere with measurements. Tracheas were stained with a 0.0125% alcian blue solution diluted in bicarbonate-buffered solution for 5 minutes at 37°C and then mounted on an inclined platform at a 30° angle, heated between 28°C and 32°C. Finally, images were acquired at 2-second intervals for 10 minutes using a Motic camera (ModelPlus 1080X) and the mucus movement was analyzed in the image sequence using Fiji, with the TrackMate-plugin and the Simple LAP tracker algorithm to identify the trajectories of alcian blue stained structures. Particles with speed zero were not included to calculate the average speed of mucus on each experiment. The mucus length was obtained using the freehand line tool from Fiji after a size calibration in µm.

### Statistical analysis

The data is presented as mean ± SEM, except for Figure 6B that is median ± SEM. Individual experiments are presented on each figure. For experiments comparing proximal vs distal from the same trachea the paired Student’s t test was used. For paired comparisons between different groups the unpaired Student’s t test was used. For comparisons between multiple groups, the ANOVA on Ranks test vs control group was used (Sigmaplot V12.3). The p<0.05 value was considered to be statistically significant. The precise information is also included in the figure legends for each data set.

## RESULTS

### Proximal and distal trachea exhibits a different electrophysiological profile

After successfully established the technique for dissection and mounting of the proximal and distal trachea of the same animal the electrophysiological studies were performed. Figure 1A, depicts the areas defined as proximal and distal trachea that were used for the electrophysiological recordings. After initial stabilization of the electrical parameters (V_TE_ and I_eq_) a protocol for assessing the effects of different blockers and agonists was applied in parallel to both tracheal sections (Figure 1B). First we observed that there was no difference in R_TE_ between proximal and distal trachea (Figure 1C), but both V_TE_ and basal I_eq_ were significantly smaller in the proximal trachea (Figure 1D and 1E), a difference that is influenced by the nearly absence of amiloride-sensitive electrogenic sodium absorption in this region (Figure 1B and F). We observed that after ENaC blockade the amiloride-non-sensitive I_eq_ was still significantly higher in the distal trachea suggesting that other transport mechanisms, besides ENaC-dependent Na^+^ absorption, are more active in this tracheal section (Figure 1G). More differences were observed in both FSK- and succinate-induced I_eq_ that were significantly higher in the proximal trachea (Figure 1B, 1H and 1I). Even though, carbachol-induced I_eq_ showed a similar magnitude in both sections (Figure 1J), the UTP-induced I_eq_ was significantly higher in the distal trachea instead (Figure 1K).

**Figure 1.**
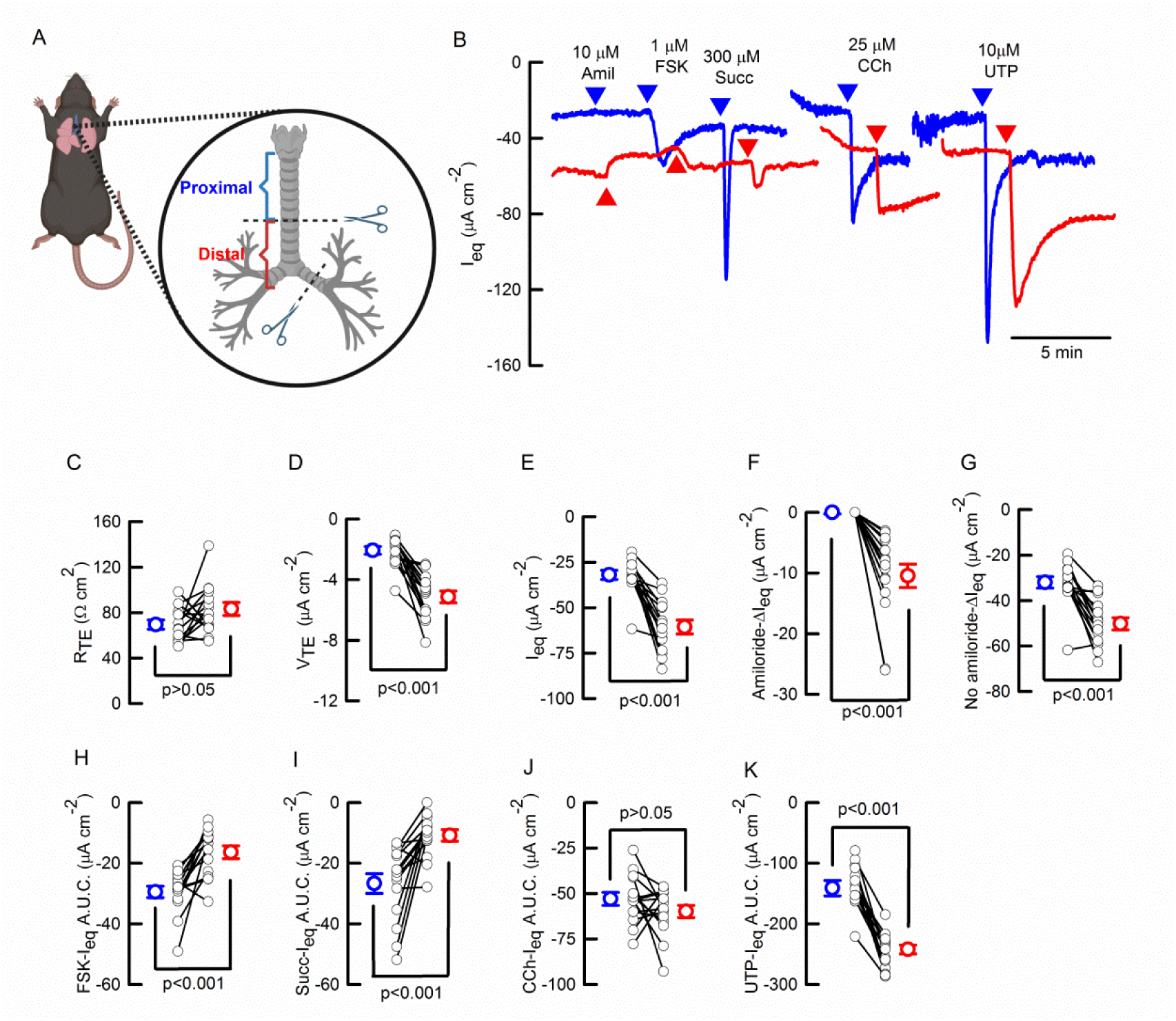
Ussing chamber electrophysiological characterization of the proximal and distal mouse trachea. (A) Depiction of proximal and distal regions of the mouse trachea used for electrophysiological measurements. (B) Representative I_eq_ traces including concentration of inhibitors and agonists. Proximal (blue) and distal (red) mouse trachea. Summary of paired determinations of (C) R_TE_, (D) V_TE_, (E) basal I_eq_, (F) amiloride-sensitive I_eq_, (G) amiloride-insensitive I_eq_, (H) forskolin-induced I_eq_, (I) Succinate-induced I_eq_, (J) carbachol-induced I_eq_ and (K) UTP-induced I_eq_. t-student paired-test; n=15. Values ±S.E.M. are included in Table 1.

**TABLE 1.**

|  | Control (5) |  | +S0859 (5) |  | +ANI9 (6) |  | *Nkcc1*<sup>+/+</sup> (5) |  | *Nkcc1*<sup>-/-</sup> (5) |  |
| --- | --- | --- | --- | --- | --- | --- | --- | --- | --- | --- |
|  | Prox | Dist | Prox | Dist | Prox | Dist | Prox | Dist | Prox | Dist |
| R<sub>TE</sub> | 70±9 | 90±14 | 76±9 | 69±12 | 77±6 | 79±6 | 98±9 | 72±7* | 107±33 | 79±3 |
| V<sub>TE</sub> | -2.3±0.7 | -5.7±1* | -1.9±0.3 | -3.0±0.6<sup>#</sup> | -1.6±0.2 | -2.0±0.2<sup>#</sup> | -2.4±0.4 | -5.7±1* | -2.8±1 | -4.2±0.8 |
| I<sub>SC</sub> | -36±7 | -60±7* | -24±2 | -43±2*<sup>#</sup> | -21±1<sup>#</sup> | -35±1*<sup>#</sup> | -29±3 | -59±10* | -25±3 | -53±8* |
| Amil-sens | 0±1 | -11±4* | -2±2 | -13±5 | -3±0.3<sup>#</sup> | -7±1 | 0±1 | -14±4* | -2±1 | 26±6*<sup>#</sup> |
| Amil-non | -36±7 | -49±4* | -23±3 | -43±2* | -19±1<sup>#</sup> | -26±2<sup>#</sup> | -29±3 | -46±7* | -23±2<sup>#</sup> | -27±3<sup>#</sup> |
| FSK | -36±4 | -17±5* | -24±2<sup>#</sup> | -16±1* | -31±2 | -23±4* | -28±3 | 23±6 | -20±1<sup>#</sup> | -12±3*<sup>#</sup> |
| Succinate | -25±5 | -10±4* | -2±2<sup>#</sup> | -2±1<sup>#</sup> | -40±6<sup>#</sup> | -11±2 | -18±8 | -12±5 | -20±5 | -8±2* |
| CCh | -62±7 | -61±9 | -49±7 | -40±7<sup>#</sup> | -46±4<sup>#</sup> | -39±6<sup>#</sup> | -60±6 | -42±5* | -32±5<sup>#</sup> | -27±3<sup>#</sup> |
| UTP | -144±26 | -244±14* | -111±18 | -139±18<sup>#</sup> | -182±20 | -202±17<sup>#</sup> | -200±17 | -220±31 | -131±31<sup>#</sup> | -109±4<sup>#</sup> |

Values are mean ± SEM. \*indicates p<0.05 for proximal versus distal on each condition. # indicates p<0.05 for proximal versus proximal or distal versus distal for control versus S0859 or *Nkcc1*<sup>+/+</sup> vs *Nkcc1*<sup>-/-</sup>. Student's t test and n indicated as (n) for each group.

### The basolateral cotransporter NBCe1 and the apical TMEM16A chloride channel contribute differentially to agonist-induced anion secretion across mouse trachea

Bicarbonate and chloride secretion have been shown to depend on NBCe1 (coded by the *Slc4a4* gene) and TMEM16A, respectively, in the mouse trachea. Thus we aimed to study their contribution to anion secretion in the proximal and distal sections using the inhibitors S0859 for NBCe1 and ANI9 for TMEM16A, which were applied in the luminal side. While S0859 induced a higher change in basal I_eq_ in the distal trachea, ANI9 reduced I_eq_ in both tracheal sections equally (Figure 2A-C). Those changes significantly reduced the magnitude of V_TE_ exclusively in the distal trachea of tissues treated with either of the blockers without affecting R_TE_ (Table 1).

**Figure 2.**
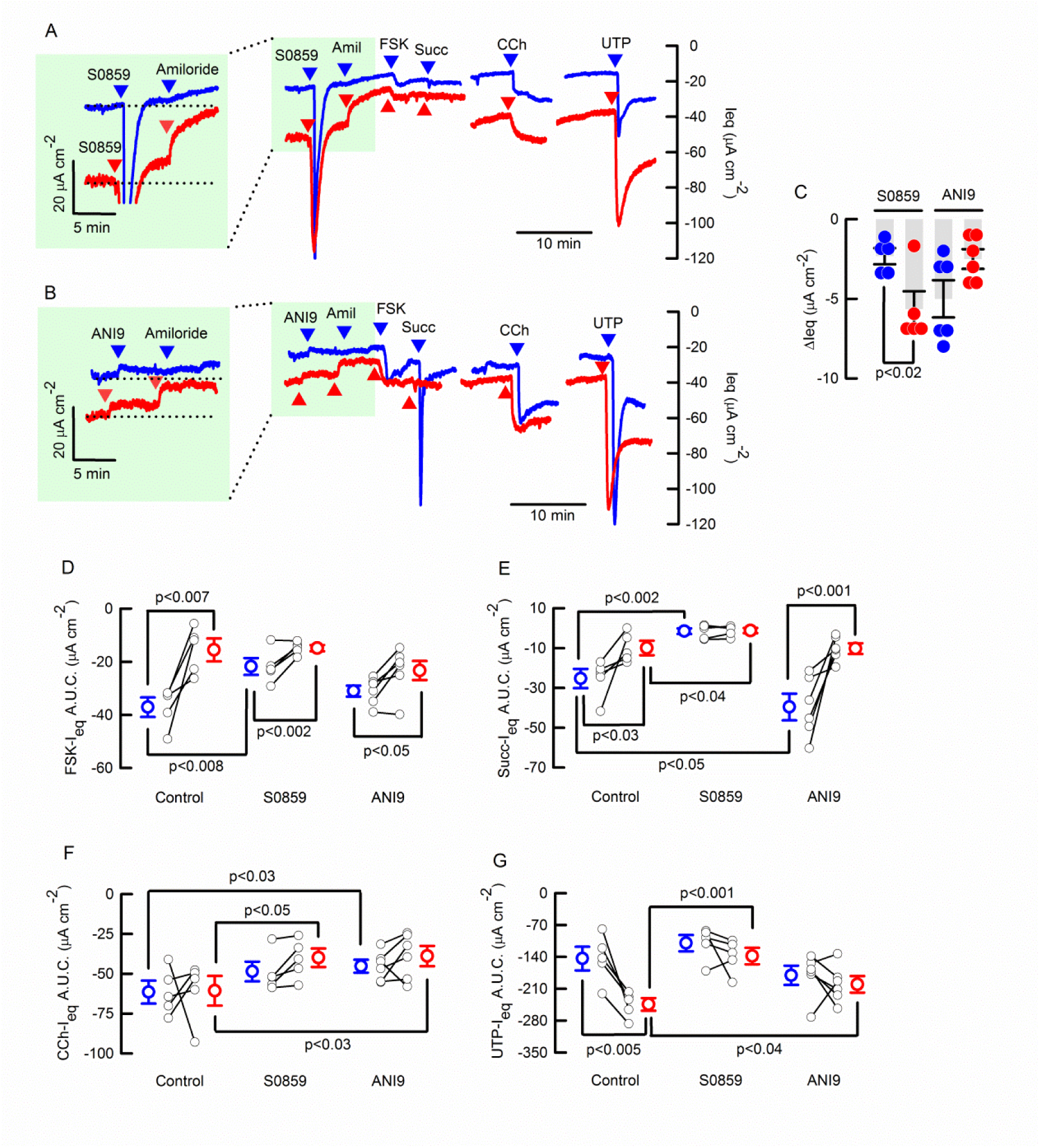
NBCe1 and TMEM16A blocking differentially affects the induced anion secretion of the proximal and distal mouse trachea. Representative traces of I_eq_ after the addition of 30 µM S0859 (A) or 30 µM ANI9 in proximal (blue line) and distal (red line) mouse tracheas. (C) Summary of the effect of S0859 and ANI9 in basal I_eq_. (D) forskolin-induced I_eq_, (E) succinate-induced I_eq_, (F) carbachol-induced I_eq_ and (G) UTP-induced I_eq_, in control, or after S0859 or ANI9 addition. Groups include n=5 control, n=5 S0859 and n=6 for ANI9; t-student paired-test for proximal versus distal on each group and t-student unpaired-test in comparisons between different groups.

Next, we observed that forskolin- and succinate-induced I_eq_ in the proximal trachea, and that succinate-, carbachol- and UTP-induced I_eq_ in the distal trachea were significantly reduced after NBCe1 inhibition with S0859 (Figure 2D-G; Table 1). Meanwhile, ANI9 showed different effects to those induced by S0859, including a significant increase in amiloride-sensitive and succinate-induced I_eq_ in the proximal section, when compared to the control group (Figure 2E & Table 1). While not affecting FSK- and succ-induced I_eq_ in the whole trachea (Figure 2D, E & Table 1), ANI9 reduced the non-sensitive amiloride I_eq_ in distal and proximal trachea, and both the CCh- and UTP-induced I_eq_ in the distal trachea exclusively (Figure 2F, G & Table 1).

### NKCC1 preferentially locates in the distal trachea and intrapulmonary airways of the mouse

We used immunolocalization to study NKCC1 distribution in the airways. As shown in Figure 3A, the proximal trachea barely expresses NKCC1 in the surface epithelium except for scarce and isolated cells with intense expression or NKCC1^high^+ (Figure 3B-C). We also observed strong NKCC1 expression in the basolateral membrane of cells from SMGs (Figure 3A-C). Patches of cells with basolateral expression of NKCC1 and NKCC1^high^+ cells were detected starting from the middle region to the most distal section of the trachea (Figure 3D-F). A stronger signal was detected in the main extrapulmonary and intrapulmonary bronchi (Supplemental Figure 1E). However, NKCC1 expression gradually decreased towards the bronchioles (Supplemental Figure 1F) and terminal bronchioles (Supplemental Figure 1G), with no detectable signal in the alveolar region. There was no signal in the *Nkcc1*^-/-^ tissue in all analyzed sections (Supplemental Figure 1B’-G’).

**Figure 3.**
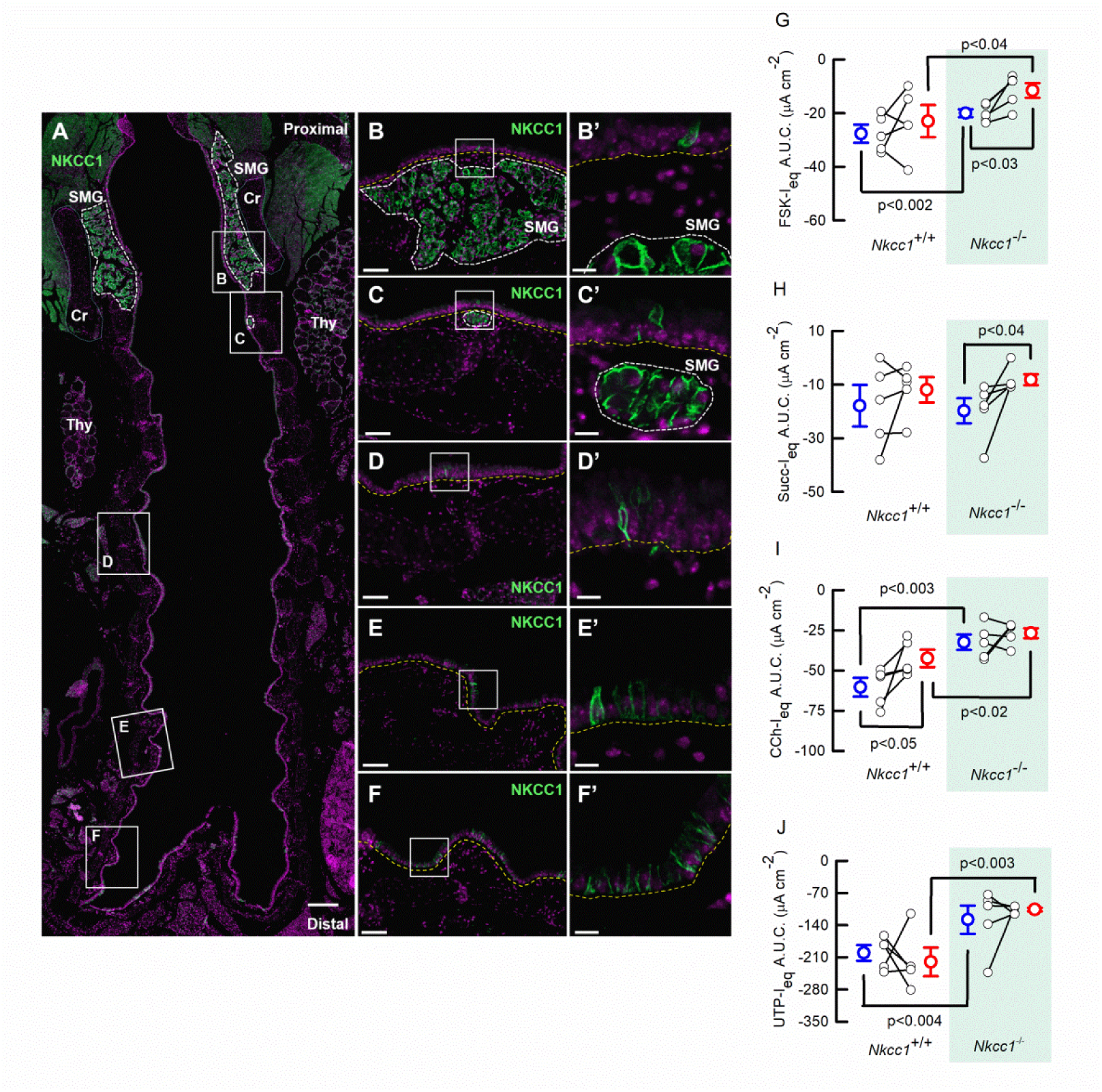
NKCC1 distribution airway epithelium and effect on secretory responses of the mouse trachea. (A) representative frontal section of the trachea from an adult wild-type (WT) mouse, showing immunostaining for the Sodium-Potassium-Chloride cotransporter 1 (NKCC1) in the surface epithelium (SAE). (B) Higher magnification of SMGs, (C) proximal, (D) medial and (E) distal regions of the tracheal and (F) and main bronchi SAE for the WT mouse. The right column (B’-F’) shows higher-magnification views of the regions indicated by the insets in panels B–F. White dashed lines outline the submucosal glands (SMGs), yellow dashed lines delineate the basal lamina of the SAE, and light blue dashed lines indicates cricoid cartilage (Cr). Nuclei are counterstained with DAPI. Scale bars: 200 µm (A), 50 µm (B–F), and 10 µm (B’–F’). Representative images of wild type (n=6). Thyroid Gland (Thy). (G) forskolin-induced I_eq_, (H) succinate-induced I_eq_, (I) carbachol-induced I_eq_ and (J) UTP-induced I_eq_, wild type *Slc12a2*^+/+^ or *Slc12a2*^-/-^ tissues; n=5 for each group.; t-student paired-test for proximal (blue dots) versus distal (red dots) on each group and t-student unpaired-test in comparisons between different groups.

To determine the contribution of the triple co-transporter NKCC1 to anion secretion, we used the *Nkcc1*-null mouse. We observed that *Nkcc1* silencing didn’t affect basal electrical parameters of the trachea, but significantly increased amiloride-sensitive sodium absorption in the distal trachea, and significantly reduced the amiloride-non-sensitive I_eq_ in both segments, evidencing the existence of basal Cl^-^ secretion in both proximal and distal trachea (Table 1). Forskolin-, carbachol- and UTP-induced I_eq_ were significantly reduced in both sections of the *Nkcc1*^-/-^ mouse trachea (Figure 3G, I and J; Table 1), but succinate-induced I_eq_ was unaffected (Figure 3H; Table1).

To ascertain if NKCC1^high^+ cells corresponded to pulmonary ionocytes we used an ASCL3 (a transcription factor expressed exclusively in ionocytes) reporter model for its identification, where the tomato protein is expressed in response to the Cre-recombinase that is under the *Ascl3* gene promoter. The co-staining determined that there were 3 different cell types NKCC1^high^+, ASCL3+ and NKCC1^high^+/ASCL3+ present in the surface airway epithelium of the trachea (Figure 4A-C).

**Figure 4.**
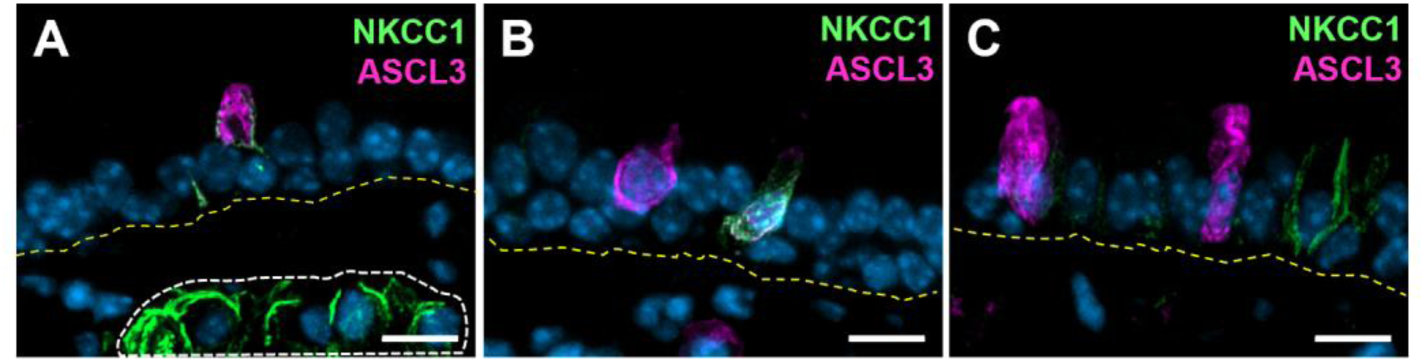
NKCC1 is expressed in some ASCL3+ cells. (A-C) Representative images from sections of the tracheal surface epithelium from an adult *Ascl3*^PGA-GCE^;*Rosa26*^tdTOMATO^ mouse, showing immunostaining for the Sodium-Potassium-Chloride cotransporter 1 (NKCC1^high^)+ cells (green) and ASCL3+ cells (Magenta). (A, B) Colocalization of NKCC1 and ASCL3 is observed in some cells (Weighted colocalization coefficients: 0.78118 and 0.99441, respectively), whereas (C) shows a lack of colocalization. Yellow dashed lines delineate the basal lamina of the tracheal epithelium, and white dashed lines outline the submucosal glands (SMGs). Nuclei are counterstained with DAPI. Scale bars 10 µm, n=3.

Comparisons of the differences between proximal and distal trachea in groups of controls for the experiments performed for the S0859 and ANI9 groups that are of the C57Bl6/J and those of the *Nkcc1*^+/+^ group that are in a hybrid background showed that differences attributable to the genetic background exists. As shown in Table 1, the C57Bl6/J mouse showed larger secretory responses to forskolin and succinate in the proximal trachea and for UTP in the distal trachea, but neither such differences are present in the *Nkcc1*^+/+^ group (Table 1). Additionally, the *Nkcc1^+/+^* tissues showed that a significant larger carbachol-induced anion secretion exists in the proximal trachea compared to the distal trachea, a phenomena that is not observed in the C57Bl6/J tissues (Table 1).

### ILs reshape the electrophysiology of the mouse trachea, mucus speed transport and shape

To study the changes induced by inflammatory signaling, we instilled mice with interleukins IL-1β, IL-17A, IL-4, or IL-13. First, we compared cytokine-induced changes in the proximal and distal tracheal regions to those observed in control BSA-instilled animals. The analysis revealed that none of the interleukins affected any measured parameter in the proximal trachea. However, in the distal trachea, IL-1β significantly reduced R_TE_ and V_TE_, while IL-17A reduced V_TE_ and basal I_eq_ (Figure 5A–C). Amiloride-sensitive Na_⁺_ absorption was reduced exclusively by IL-17A, whereas amiloride-insensitive I_eq_ remained unchanged across all treatments (Figure 5D & E). Forskolin-, succinate-, and UTP-induced I_eq_ were increased by IL-4, whereas carbachol- and UTP-dependent I_eq_ were significantly reduced by IL-17A. A summary of these changes is presented in Figure 5J.

**Figure 5.**
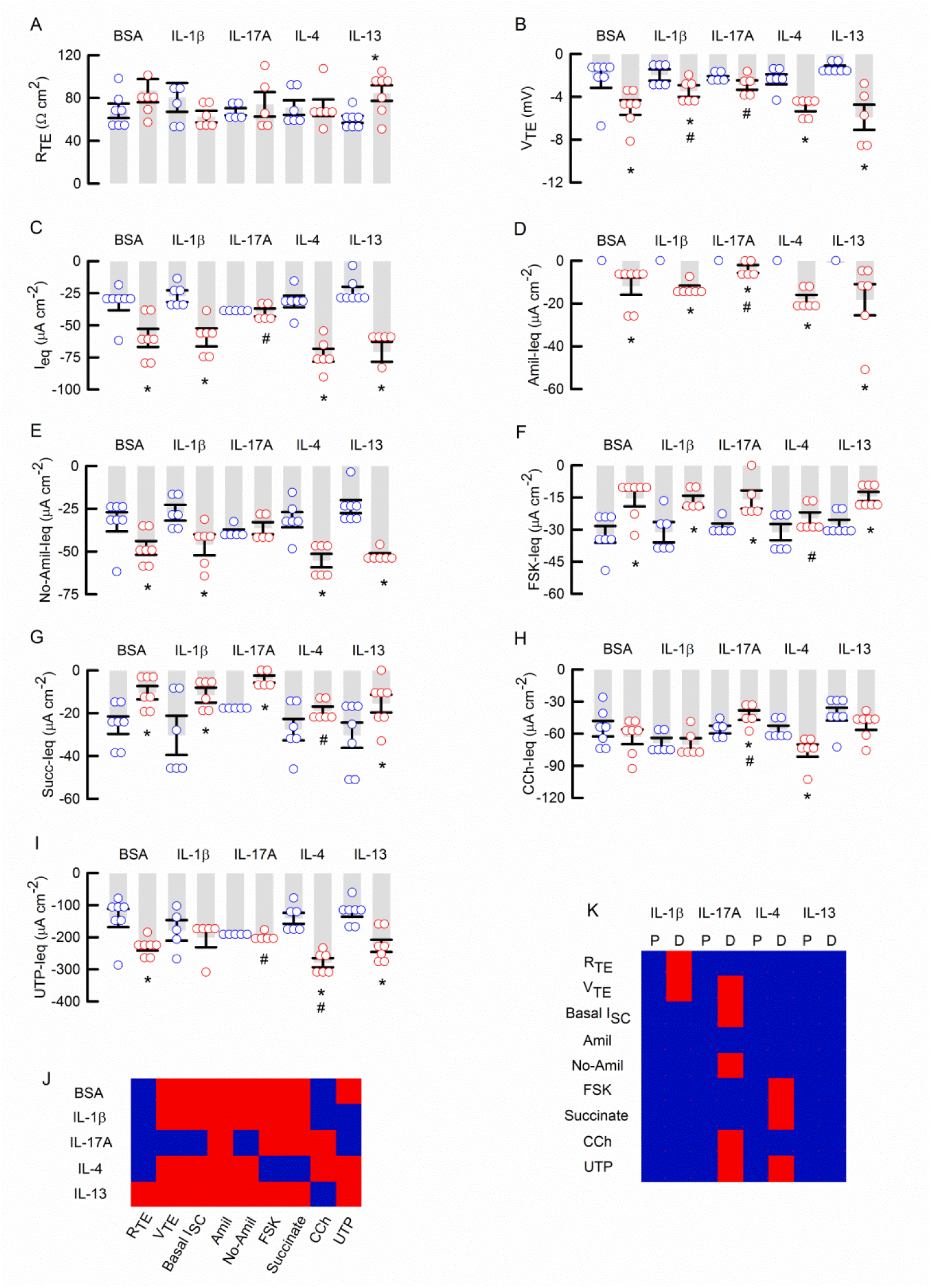
The effect of interleukins on the electrophysiological properties of proximal and distal mouse trachea. The figure include the changes in proximal (blue circles) and distal (red circles) trachea induced by 30ng of either IL1B (n=6), IL17A (n=5), IL-4 (n=6) or IL13 (n=7) and controls (BSA: n=6) in (A) R_TE_, (B) V_TE_, (C) Basal-I_eq_, (D) amiloride-sensitive I_eq_, (E) amiloride-insensitive I_eq_, (F) forskolin-induced I_eq_, (G) Succinate-induced I_eq_, (H) carbachol-induced I_eq_ and (I) UTP-induced I_eq_. * indicates p<0.05 proximal vs distal. # indicates p<0.05 vs BSA. Panel (J) summarizes distal vs proximal statistical differences and panel (K) summarizes ILs effect vs BSA blue for p>0.05 and red for p<0.05. Data is also available at Table 2.

**TABLE 2.**
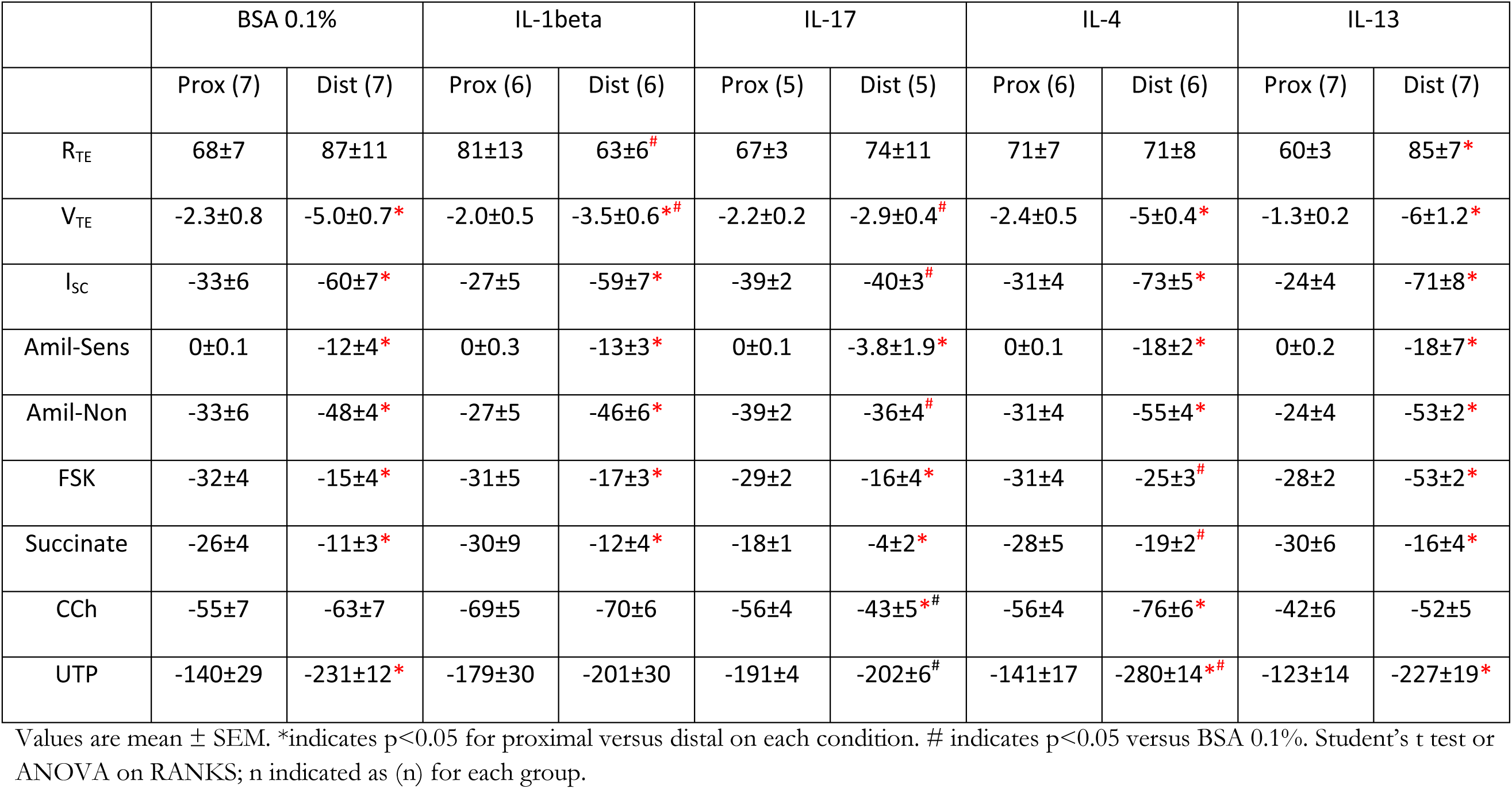

|  | BSA 0.1% |  | IL-1beta |  | IL-17 |  | IL-4 |  | IL-13 |  |
| --- | --- | --- | --- | --- | --- | --- | --- | --- | --- | --- |
|  | Prox (7) | Dist (7) | Prox (6) | Dist (6) | Prox (5) | Dist (5) | Prox (6) | Dist (6) | Prox (7) | Dist (7) |
| R<sub>TE</sub> | 68±7 | 87±11 | 81±13 | 63±6<sup>#</sup> | 67±3 | 74±11 | 71±7 | 71±8 | 60±3 | 85±7<sup>*</sup> |
| V<sub>TE</sub> | -2.3±0.8 | -5.0±0.7<sup>*</sup> | -2.0±0.5 | -3.5±0.6<sup>*#</sup> | -2.2±0.2 | -2.9±0.4<sup>#</sup> | -2.4±0.5 | -5±0.4<sup>*</sup> | -1.3±0.2 | -6±1.2<sup>*</sup> |
| I<sub>SC</sub> | -33±6 | -60±7<sup>*</sup> | -27±5 | -59±7<sup>*</sup> | -39±2 | -40±3<sup>#</sup> | -31±4 | -73±5<sup>*</sup> | -24±4 | -71±8<sup>*</sup> |
| Amil-Sens | 0±0.1 | -12±4<sup>*</sup> | 0±0.3 | -13±3<sup>*</sup> | 0±0.1 | -3.8±1.9<sup>*</sup> | 0±0.1 | -18±2<sup>*</sup> | 0±0.2 | -18±7<sup>*</sup> |
| Amil-Non | -33±6 | -48±4<sup>*</sup> | -27±5 | -46±6<sup>*</sup> | -39±2 | -36±4<sup>#</sup> | -31±4 | -55±4<sup>*</sup> | -24±4 | -53±2<sup>*</sup> |
| FSK | -32±4 | -15±4<sup>*</sup> | -31±5 | -17±3<sup>*</sup> | -29±2 | -16±4<sup>*</sup> | -31±4 | -25±3<sup>#</sup> | -28±2 | -53±2<sup>*</sup> |
| Succinate | -26±4 | -11±3<sup>*</sup> | -30±9 | -12±4<sup>*</sup> | -18±1 | -4±2<sup>*</sup> | -28±5 | -19±2<sup>#</sup> | -30±6 | -16±4<sup>*</sup> |
| CCh | -55±7 | -63±7 | -69±5 | -70±6 | -56±4 | -43±5<sup>*#</sup> | -56±4 | -76±6<sup>*</sup> | -42±6 | -52±5 |
| UTP | -140±29 | -231±12<sup>*</sup> | -179±30 | -201±30 | -191±4 | -202±6<sup>#</sup> | -141±17 | -280±14<sup>*#</sup> | -123±14 | -227±19<sup>*</sup> |

Values are mean ± SEM. \*indicates p<0.05 for proximal versus distal on each condition. # indicates p<0.05 versus BSA 0.1%. Student's t test or ANOVA on RANKS; n indicated as (n) for each group.

Next, we assessed whether the effects of interleukins altered the proximal-distal differences compared to those observed in BSA-instilled controls. The analysis showed that R_TE_ exhibited a significant proximal-distal difference after IL-13 treatment (Figure 5A). Due to the significant reduction in V_TE_ and basal I_eq_ in the distal trachea, the proximal-distal difference was lost in response to IL-17A treatment (Figure 5B & C). IL-17A also affected amiloride-insensitive I_eq_ (Figure 5E), while amiloride-sensitive I_eq_ remained unchanged across all treatments (Figure 5D).

The significant increase in FSK- and succinate-induced I_eq_ in the distal trachea after IL-4 treatment eliminated the statistical difference between regions (Figure 5F & G). Carbachol-induced I_eq_ was differentially affected by IL-17A and IL-4: while IL-17A reduced it, IL-4 significantly increased I_eq_ in the distal trachea (Figure 5H). Finally, IL-1β and IL-17A altered UTP-induced I_eq_, leading to a loss of statistical significance between proximal and distal trachea (Figure 5I). A summary of these statistical differences is shown in Figure 5K.

As observed in Fig 6A the instillation of ILs induced a significant increase in basal mucus speed transport in the IL-13 instilled animals exclusively. But an increase in the length of mucus structures was observed across all groups of ILs instilled animals (Figure 6B). BSA 0.1% instilled animals exhibited mucus structures in a range of 75-300 µm length in the form of small clouds or flocks and very few threads with a median length value of 145 µm. The effect of ILs significantly increased the median length to 296 µm (IL-1 β), 305 µm (IL-17A), 210 µm (IL-4) and 236 µm (IL-13). All ILs induced the apparition of bigger threads between the 400-1500 µm in length. Only IL-4 induced the apparition of clouds of mucus of 400 µm in length. Examples of the mucus structures observed are shown in Figure 6C.

**Figure 6.**
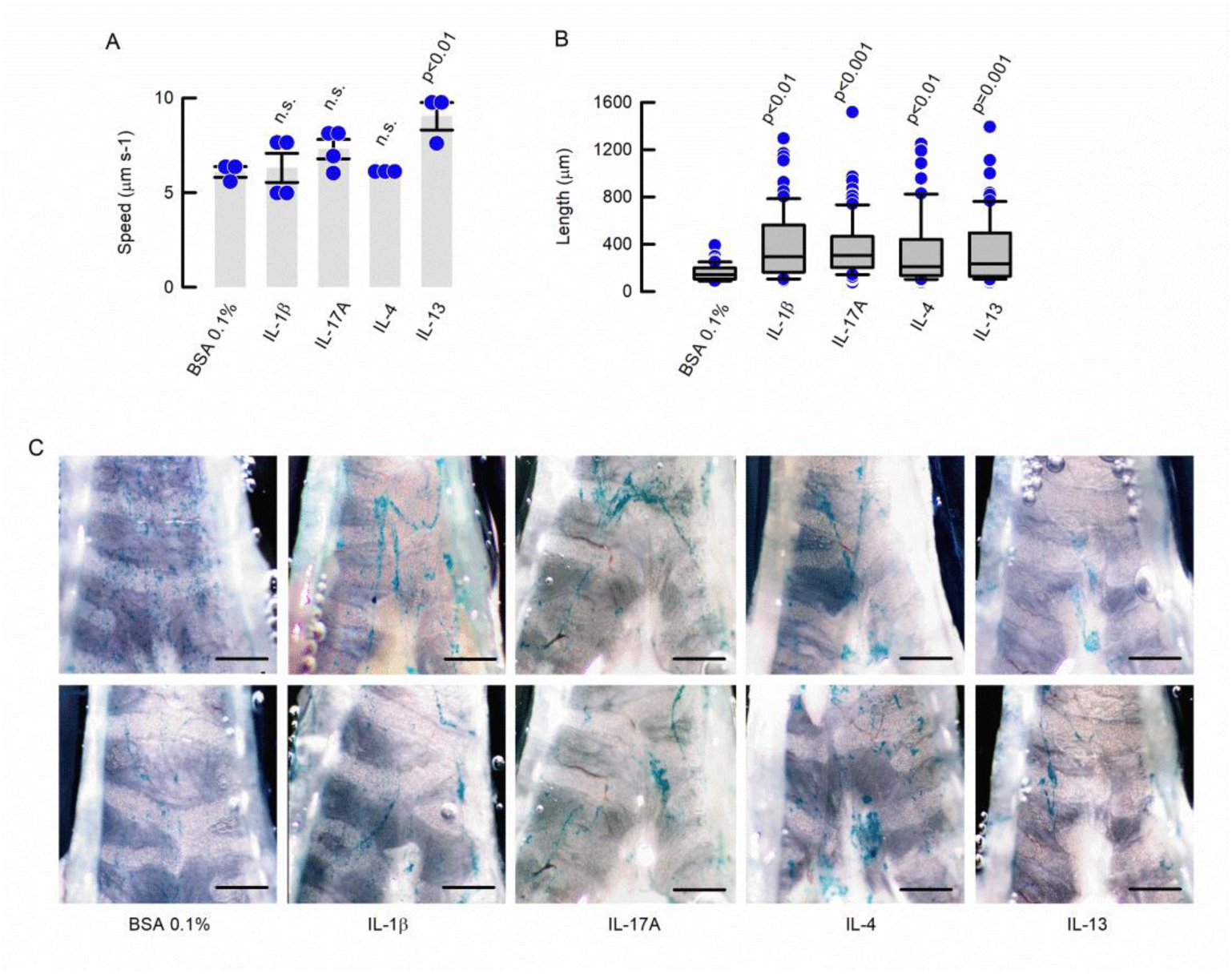
ILs increase mucus speed and change mucus structures in the mouse airways. (A) Summary of mucus speed (BSA 0.1%, IL-4 and IL-13 n=3 each, IL-1B and IL-17 n=4 each; ANOVA on Ranks) and (B) length of mucus after ILs instillation in the mouse trachea (BSA 0.1% n=64, IL1β n=78, IL-17A n=80, IL-4 n=60 and IL-13 n=61, taken from the same experiments in A; ANOVA on Ranks). (C) Representative images of mucus structures stained with alcian blue, for 2 different animals for each group. Bar indicates 1mm.

## DISCUSSION

The main results of the present study demonstrate: 1) for the first time, and after minimal modifications, that electrophysiological recordings can be obtained from both the proximal and distal trachea; 2) Ion transport modulation exhibits regional differences; and 3) cytokines can independently modulate ion transport mechanisms in both mouse tracheal regions.

Since most tissue sliders expose only a small area of dissected tracheal tissue, Ussing chamber experiments likely record I_eq_ from either proximal middle or distal epithelial regions, leading to high variability and reduced reproducibility. For instance, the striking difference in ENaC activity between the proximal and distal trachea may explain the reported variability and challenges in measuring ENaC-dependent I_eq_ (Grubb *et al*., 2001; Gianotti *et al*., 2016; Vega *et al*., 2020; Philp *et al*., 2022). Furthermore, we found that NKCC1 absence induced a twofold increase in ENaC-mediated sodium absorption in the distal trachea, consistent with the higher density of NKCC1+ cells in this region suggesting that intracellular Na_⁺_ depletion due to NKCC1 absence enhances the gradient for Na_⁺_ absorption. This difference, however, was not observed in whole-trachea electrophysiological recordings (Grubb *et al*., 2001). Our data also indicate that the distal trachea maintains a significantly more negative V_TE_ and higher basal I_eq_, sustained by greater amiloride-sensitive Na_⁺_ absorption, basal NBCe1- and TMEM16A-depdendent anion secretion in this region. Although NKCC1-dependent basal chloride secretion does not significantly contribute to V_TE_ or basal I_eq_, its absence significantly reduced amiloride-insensitive I_eq_ in both proximal and distal epithelium, pointing to a minor role in basal chloride secretion across the entire trachea, that also might explain the absence of a muco-obstructive phenotype after genetic silencing in the mice (Grubb *et al*., 2001).

Forskolin is widely used to activate cAMP-dependent anion secretion in various tissues, including the mouse trachea (Millar-Büchner *et al*., 2016; Vega *et al*., 2020). We observed that FSK-induced anion secretion was higher in the proximal trachea and significantly reduced by S0859 or after silencing of NKCC1, but not by ANI9 ruling out a relevant role of TMEM16A in cAMP-induced secretion. Notably, the dependence of NKCC1 was also observed using cultured mouse tracheal cells but not in experiments using the whole trachea of *Slc12a2*^-/-^ mice after FSK stimulation (Flagella *et al*., 1999; Grubb *et al*., 2001). Although NKCC1^high^+ cells are scarce in the proximal surface airway epithelium, the co-localization with ASCL3 suggest that some NKCC1^high^+ cells correspond to CFTR-rich pulmonary ionocytes that together with NKCC1+ cells in submucosal glands (SMGs), could be behind the NKCC1-dependent FSK-induced secretion in the proximal trachea. The last is supported by previous observations of FSK-stimulated CFTR-mediated secretion in SMGs (Ianowski *et al*., 2007). The combined chloride (NKCC1-dependent) and bicarbonate (NBCe1-dependent) secretion triggered by FSK may play a key role in SMG function, similar to what has been proposed for mucus expulsion in intestinal crypts in accordance to the relevant role carried by cAMP-activated CFTR but not Ca^2+^-activated TMEM16A channels (Vega *et al*., 2019; Dolan *et al*., 2022). This secretory process is crucial because chloride secretion facilitates mucus expulsion from SMGs, while bicarbonate secretion is critical for mucus maturation, goblet cell release, and proper mucociliary clearance (MCC) in the airway epithelium (Chen *et al*., 2010; Gorrieri *et al*., 2016; Saint-Criq *et al*., 2022).

We observed that succinate-induced anion secretion is significantly higher in the proximal trachea. Although the distribution of the succinate receptor SUCNR1 has not been determined in the mouse and human trachea, an alternative explanation to the observed regional difference in response to the addition of succinate might stems from variations in the number of anion-secreting cells activated downstream of brush cell-mediated acetylcholine release (Apablaza *et al*., 2025). Even though, acetylcholine released by brush cells exerts its paracrine effect via muscarinic receptor 3 (M3R)-positive cells, no proximal-distal difference in anion secretion stimulated with CCh (another M3R agonist) was observed. This rules out uneven M3R expression across the proximal-distal axis as the cause of the differential succinate-induced I_eq_, implying that SUCNR1 distribution or other, yet unidentified factors underlie this disparity (Perniss *et al*., 2023; Billipp *et al*., 2024; Apablaza *et al*., 2025). Consistent with our prior findings that succinate-induced I_eq_ is primarily bicarbonate-driven, we observed near-complete inhibition upon blocking NBCe1 with S0859, but was not affected in the *Nkcc1*^-/-^ mouse (Apablaza *et al*., 2025). Though our previous studies on whole tracheas also showed that ANI9 did not affect succinate-induced anion secretion, ANI9 induced an increase in succinate-induced I_eq_ in the proximal trachea. This intriguing effect cannot be explained by the present evidence.

Both carbachol- and UTP-induced I_eq_ exhibited the same dependence on NBCe1 only in the distal trachea. Our data also point towards an important role for TMEM16A and NKCC1 in the CCh-induced chloride secretion in both tracheal sections. These results indicate that the channel and the co-transporter might co-express in the same cells that respond to direct MR3 or P2Y2 stimulation. Previous studies using whole trachea preparations similarly reported that UTP-induced I_eq_ was significantly reduced in *Slc12a2*_⁻_ ^/^_⁻_ tissues or upon S0859 inhibition (Grubb *et al*., 2001; Saint-Criq *et al*., 2022). Interestingly, CCh-induced secretion from submucosal glands (SMGs) was shown to be markedly diminished by the NKCC1 inhibitor bumetanide (Ianowski *et al*., 2007), suggesting that NKCC1 activity in SMGs contributes to the CCh-induced I_eq_ observed in our experiments, which is supported by the intense NKCC1 staining detected in SMGs. Additionally, our approach also enabled the detection of differences in agonist-induced anion secretion across mouse strains, a phenomenon previously documented in the colon (Flores *et al*., 2010).

Cytokines have been extensively used to study fluid absorption and secretion in human bronchial epithelial cells (HBECs), leading to key discoveries such as the identification of the Ca²_⁺_ -activated chloride channel TMEM16A (Caputo *et al*., 2008), however, electrophysiological experiments using cytokines in the mouse airways remain limited (Anagnostopoulou *et al*., 2010; Philp *et al*., 2022). In our experiments, none of the tested interleukins altered agonist-induced anion secretion in the proximal trachea. But, most notably, in the distal trachea IL-1β and IL-17A induced changes ascribed to an anti-secretory phenotype, while IL-4 was pro-secretory, and IL-13 produced no significant changes. Whether this regional specificity stems from preferential IL-receptor expression in the distal trachea or would emerge in the proximal trachea at higher cytokine concentrations remains unclear. Compared to our prior work with IL-4, we now detected increased FSK-induced anion secretion in the distal trachea, a finding obscured in whole-trachea experiments (Philp *et al*., 2022). A similar pro-secretory effect of IL-4 has been observed in in human bronchial epithelial cells (HBECs) with enhanced FSK- and UTP-dependent anion and secretion that promotes bicarbonate secretion critical for mucus maturation and release (Gorrieri *et al*., 2016) (Simões *et al*., 2021). In contrast to our mouse data, IL-1β in HBECs reduced Na_⁺_ absorption while increasing cAMP- and UTP-induced anion secretion (Gray *et al*., 2004), though a subsequent study where only the Na_⁺_ absorption reduction was observed (Simões *et al*., 2021). IL-17A induced a significant decrease in basal electrical parameters (V_TE_ and I_eq_), likely due to reduced ENaC-mediated Na_⁺_ absorption. It also markedly suppressed carbachol- and UTP-induced anion secretion in the distal trachea. IL-1β reduced R_TE_ and V_TE_ magnitude compared to controls but did not altered other parameters that might explain this effect. In contrast to our murine findings, IL-17 treatment in human bronchial epithelial cells (HBECs) increased forskolin-induced bicarbonate secretion via a CFTR-dependent mechanism, without affecting amiloride-sensitive I_eq_ (Kreindler *et al*., 2009). Some of those differences are attributable to culture conditions, species related, the use of fresh tissues versus cultured cells and doses and time of incubation with ILs.

Since changes produced by ILs concentrated in the distal region of the trachea, the experiments of mucus transport were recorded in that section. We observed that only IL-13 increased mucus transport speed in basal conditions. Paradoxically, IL-13 didn’t induce changes in the basal electrical parameters in the distal trachea. Conversely, the important reductions in V_TE_ and I_eq_ after IL-1β or IL17A instillation, predicted a decrease in mucus transport but the results shown no differences. This apparent contradiction might be explained by the fact that we exposed the animals to the ILs for a short period of time and that a more sustained inflammatory signaling might produce negative changes in the mucus transport machinery compromising the expression of channels and transporters or altering the number of ciliated cells like shown for IL-4 in HBECs (Simões *et al*., 2021). Another explanation is that a modification in the composition of mucins will favour mucus transport. The last option is supported by the observed changes in mucus transport speed after silencing of either MUC5AC or MUC5B in the mouse and the increase in goblet cells and mucus release by effect of ILs (Gray *et al*., 2004; Anagnostopoulou *et al*., 2010; Gorrieri *et al*., 2016; Fakih *et al*., 2020; Simões *et al*., 2021). Even though, mucus speed transport can be altered by agonists of anion secretion like CCh or succinate we left that aspect unexplored in the present work (Ermund *et al*., 2018; Ehre *et al*., 2023; Apablaza *et al*., 2025).

In summary, we demonstrated that the proximal and distal trachea compartmentalized functions controlling fluid movement and responses to inflammatory cues. The proximal region exhibit greater FSK-induced secretion, probably related to SMGs activity, and succinate-induced secretion that participates in immune surveillance at the entry site of environmental air. The resilience of the proximal trachea to ILs-induced changes suggests the necessity that those functions remain constant for proper lung health. The bigger anion secretion and the presence of sodium absorption observed in the distal trachea favour MCC, and are critical for mucus transport to the proximal section. The distal trachea rapidly respond to ILs signals to adapt to increased mucus release from distal airways, otherwise accumulates in the main bronchi preferentially (Mall *et al*., 2004; Vega *et al*., 2020). The conclusions derived from our work are limited due to absence of a detailed detection of channels like ENaC and CFTR, transporters like NBCe1 and receptors like SUCNR1, MR3 or P2Y2, whose co-expression define different secretory cell types along the proximal-distal axis and are behind the functional observations presented. Changes in such cell types in the distal trachea might also be responsible of observed effects of ILs instillation, as widely documented in HBECs. Although ILs effects have historically been viewed as pathological, they are also a component of the normal immune responses. Recent insights into these mechanisms in cultured HBECs have revealed positive synergistic effects of ILs with therapies for cystic fibrosis. In some cases, this synergy is enhanced by an increased population of cells expressing proteins that participate in anion secretion (Rehman *et al*., 2021; Borrelli *et al*., 2025). Identifying the specific airway regions and cell types that respond to ILs could enable precise targeted therapies for muco-obstructive diseases. Furthermore, our findings provide a framework for testing novel synergistic pharmacological interactions with cytokines *in vivo*.

## Acknowledgements

This work was supported by FONDECYT Regular 1221257, ANID Anillo de Tecnología ACT250073 NanoAir and The Cystic Fibrosis Trust U.K. (C.A.F.); ANID PhD Fellowship No. 21250795 (T.A.), FONDECYT Postdoctoral 3220672 (S.V.) & Southern Bioimaging Platform (SBIP)-USS (FONDEQUIP-EQY240004).

## Author contributions

**Conceptualization:** C.A. Flores; **Data curation:** T. Apablaza, S. Villanueva, A. Olave-Ruiz, A. Guequen & C.A. Flores; **Formal Analysis:** T. Apablaza, S. Villanueva, A. Guequen & C.A. Flores; **Funding aquisition:** S. Villanueva & C.A. Flores; **Investigation:** T. Apablaza, S. Villanueva, A. Olave-Ruiz, A. Guequen, M.A. Catalán & C.A. Flores; **Writting – original draft:** C.A. Flores. **Writting – review & editing:** M.A. Catalán & C.A. Flores.

---

**Supplemental Figure 1.**
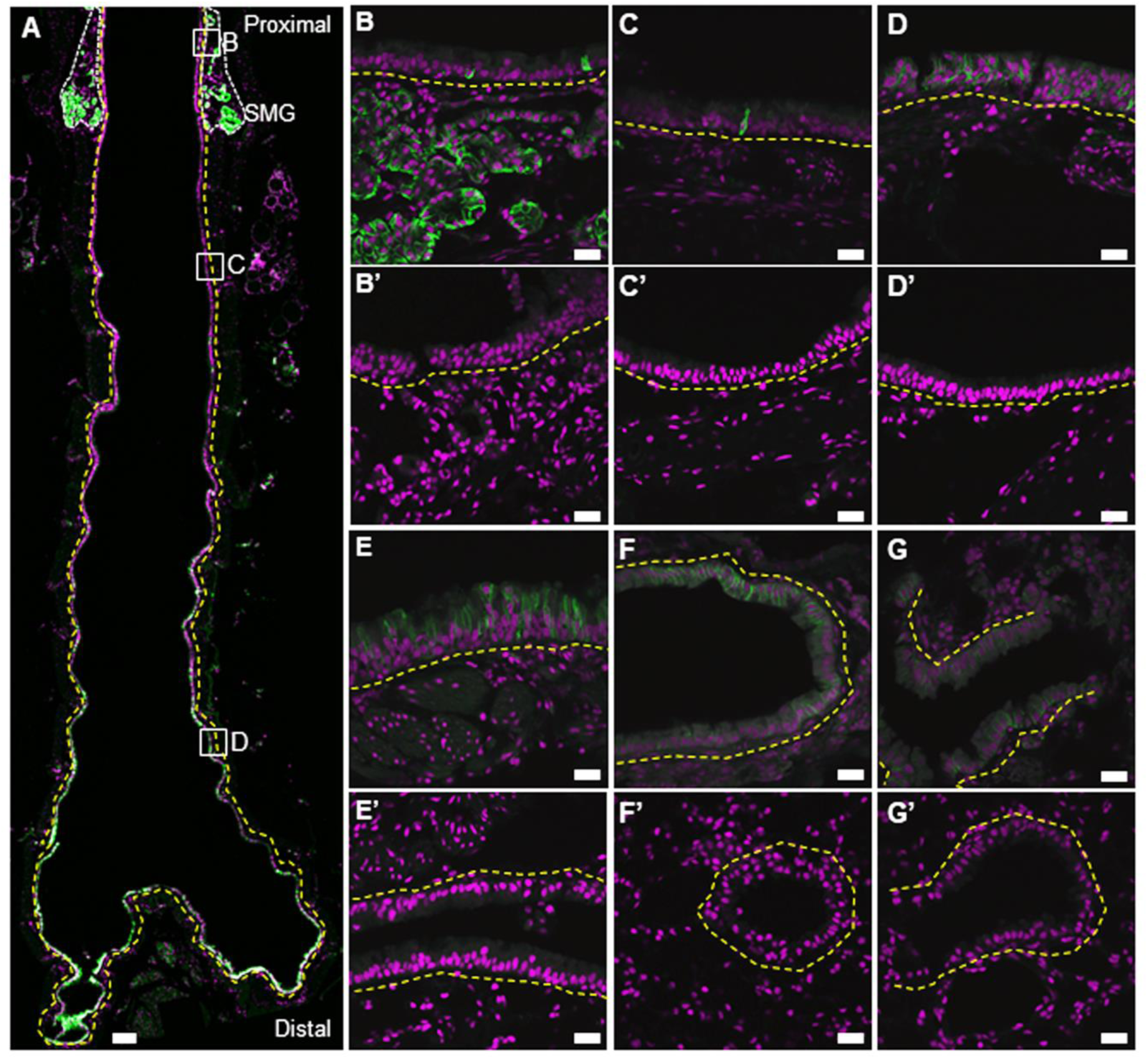
Validation of the anti-NKCC1 antibody in the mouse airways. (A) representative frontal section of the trachea from an adult wild-type (WT) mouse, showing immunostaining for the Sodium-Potassium-Chloride cotransporter 1 (NKCC1) in the surface airway epithelium (SAE). (B) Higher magnification of SMGs, (C) proximal and (D) distal regions of the tracheal SAE for the wild type (A-D) and *Slc12a2*^-/-^ (B’-D’). Representative images of NKCC1+ cells in the (E) main bronchi, (F) bronchi and (G) terminal bronchi, for wild type (E-G) and *Slc12a2*^-/-^ (E’-G’). White dashed lines outline the submucosal glands (SMGs), and yellow dashed lines delineate the basal lamina of the SAE. Nuclei are counterstained with propidium iodide. Scale bars: 100 µm (A), 20 µm (B–G). Representative images of wild type (n=4) and *Slc12a2*^-/-^ (n=3).

